# Commensal protists in reptiles display flexible host range and adaptation to ectothermic hosts

**DOI:** 10.1101/2023.05.25.542353

**Authors:** Elias R. Gerrick, Leila B. DeSchepper, Claire M. Mechler, Lydia-Marie Joubert, Freeland Dunker, Timothy J. Colston, Michael R. Howitt

## Abstract

Parabasalid protists recently emerged as keystone members of the mammalian microbiota with important effects on their host’s health. However, the prevalence and diversity of parabasalids in wild reptiles and the consequences of captivity and other environmental factors on these symbiotic protists are unknown. Reptiles are ectothermic, and their microbiomes are subject to temperature fluctuations, such as those driven by climate change. Thus, conservation efforts for threatened reptile species may benefit from understanding how shifts in temperature and captive breeding influence the microbiota, including parabasalids, to impact host fitness and disease susceptibility. Here, we surveyed intestinal parabasalids in a cohort of wild reptiles across three continents and compared these to captive animals. Reptiles harbor surprisingly few species of parabasalids compared to mammals, but these protists exhibited a flexible host-range, suggesting specific adaptations to reptilian social structures and microbiota transmission. Furthermore, reptile-associated parabasalids are adapted to wide temperature ranges, although colder temperatures significantly altered the protist transcriptomes, with increased expression of genes associated with detrimental interactions with the host. Our findings establish that parabasalids are widely distributed in the microbiota of wild and captive reptiles and highlight how these protists respond to temperature swings encountered in their ectothermic hosts.

## Introduction

The intestinal microbiota critically shapes host health through its wide-ranging influence on physiology, including metabolic and immune function. These microbes are acquired from the environment and often reflect kinship, social structure, and diet in many mammals. However, reptiles frequently perform limited or no parental care and are ectothermic, thus their microbiome is highly susceptible to reconfiguration by environmental changes^1,2^. Climate change has an outsized negative impact on reptile species density and diversity, and the microbiomes of these ectotherms are exposed to changing global temperatures^3,4^. Compounding this issue, captivity has strong impacts on reptile microbiome diversity and composition, which should be considered in conservation efforts involving captive breeding programs^5,6^. Therefore, understanding animal microbiomes is critical for developing conservation strategies for endangered or at-risk species.

Studies on the effects of temperature and captivity on reptile microbiotas have focused on bacteria, but how these environmental factors influence symbiotic eukaryotes is unknown^6^. Protists were long ignored as part of the microbiota, but recent work highlighted their profound effects on host health^7–10^. In particular, symbiotic protists in the phylum *Parabasalia* are found widely in animals from insects to humans, where they contribute to nutrient acquisition^11^, immunomodulation^7,8^, and protection from infections^8^. A limited number of studies identified parabasalids in reptiles, but the distribution and biology of these protists is poorly understood^12–14^. In this work, we catalog the parabasalid species in the gut microbiomes of a diverse array of captive and wild reptiles. We then directly interrogate the effect of temperature on these protists and find that reptile-associated parabasalids, unlike human-associated parabasalids, adapt to temperature fluctuations by inducing massive transcriptomic remodeling with predicted effects on host-microbe interactions.

## Results

To better characterize the diversity and prevalence of parabasalids in reptiles, we collected cloacal swabs from 33 wild reptiles across 3 continents (Figure 1A, Table 1). In addition, we collected stool samples from 9 captive reptiles to investigate the effects of captivity on parabasalid colonization and diversity. In total, this cohort represents 38 different reptile species. We designed pan-parabasalid primers that amplify the ribosomal internally transcribed spacer (ITS) to identify and phylogenetically align commensal parabasalids from these reptile samples. We found that 7 (21%) of the wild reptile cloacal swabs were positive for parabasalids, and 7 (78%) of the captive reptile stool samples were positive (Figure 1A). The higher proportion of positivity in captive reptiles likely reflects the difference in sampling method, as cloacal swabs provide substantially less biomass and thus are more likely to produce false negatives.

**Figure 1:**
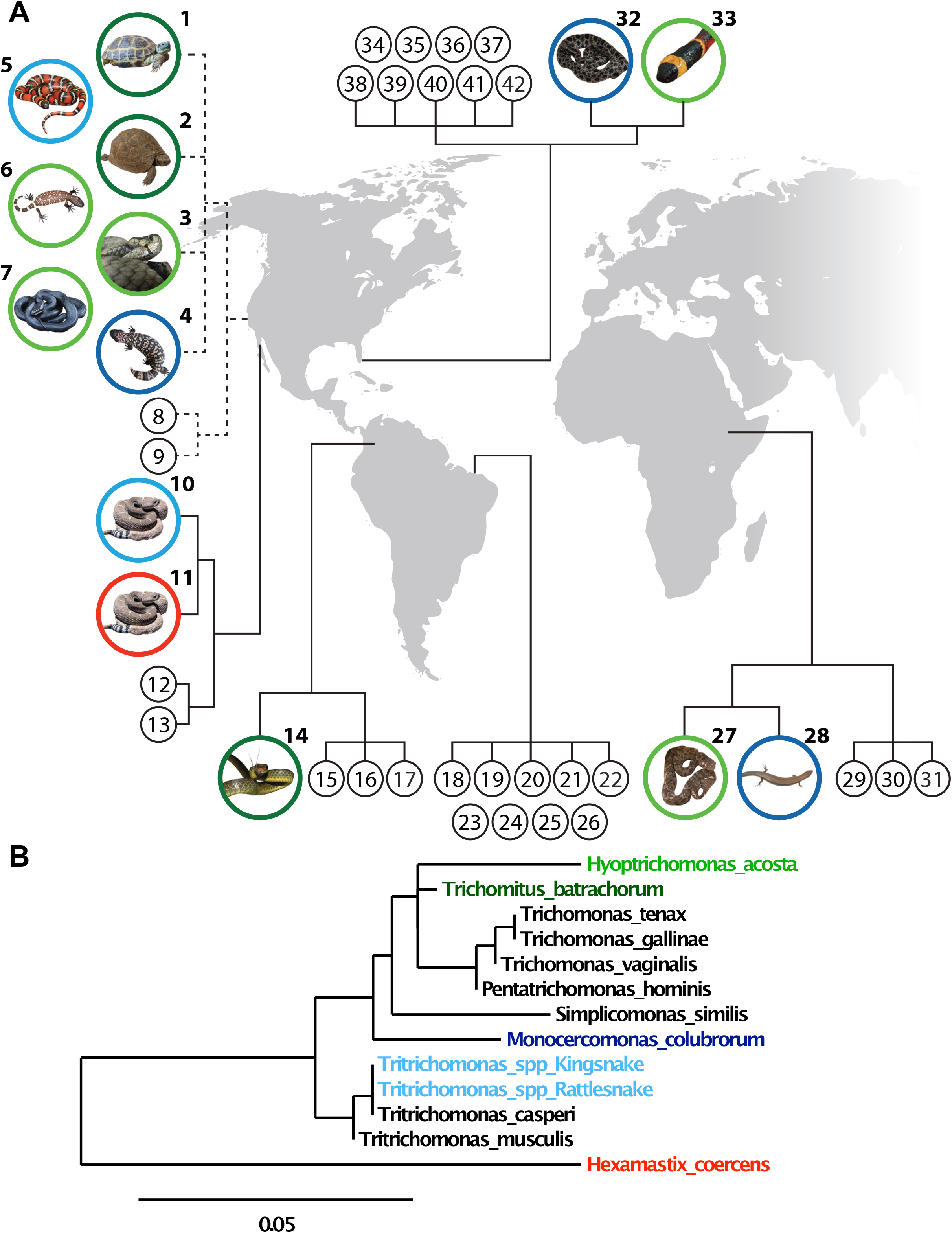
Parabasalids display limited diversity and flexible host ranges in wild and captive reptiles. (A) Locations and associated parabasalid protists of reptiles profiled in this study. Black numbered circles represent reptiles for which no parabasalid was detected. Colored circles indicate reptiles for which a parabasalid was identified in the sample; the image inside the circle depicts the reptile species, and the circle color indicates the parabasalid species identified in the reptile. Green circles indicate class *Trichomitea* (dark green, *T. batrachorum*; light green, *H. acosta*). Blue circles indicate class *Tritrichomonas* (dark blue, *M. colubrorum*; light blue, *Tritrichomonas spp*.). Red circles indicate *H. coercans*. The region each reptile was sampled is indicated on the global map. Dashed lines denote captive reptiles, solid lines denote wild reptiles sampled in their native habitat. (B) Phylogenetic tree of parabasalids identified in this study, as well as additional parabasalids of other animals including rodents and humans. Parabasalids identified in reptile samples are color-coded to match the colors in Figure 1A. The two novel *Tritrichomonas* species identified in rodent-eating snakes are labelled with the snake in which they were identified.

**Table 1.**

| Reptile number | Reptile species | Common name | Trichomonad present | Stool/Cloacal swab | Wild/Captive | Reptile location |
| --- | --- | --- | --- | --- | --- | --- |
| 1 | Testudo horsfieldii | Russian tortoise | *Trichomitus batrachorum* | Stool | Captive | California, USA |
| 2 | Testudo graeca | Greek tortoise | *Trichomitus batrachorum* | Stool | Captive | California, USA |
| 3 | Crotalus horridus | Timber Rattlesnake | *Hypotrichomonas acosta* | Stool | Captive | California, USA |
| 4 | Heloderma suspectum | Gila monster | *Monocercomonas colubrorum* | Stool | Captive | California, USA |
| 5 | Lampropeltis zonata | Mountain Kingsnake | *Tritrichomonas spp.* | Stool | Captive | California, USA |
| 6 | Heloderma horridum | Beaded Lizards | *Hypotrichomonas acosta* | Stool | Captive | California, USA |
| 7 | Pantherophis obsoletus | Western Ratsnake | *Hypotrichomonas acosta* | Stool | Captive | California, USA |
| 8 | Pituophis catenifer | Gopher snake |  | Stool | Captive | California, USA |
| 9 | Elgaria multicarinata | Alligator lizard/Lizandro |  | Stool | Captive | California, USA |
| 10 | Crotalus ruber | red rattlesnake | *Tritrichomonas spp.* | Cloacal swab | Wild | Baja, Mexico |
| 11 | Crotalus ruber | red rattlesnake | *Hexamastix coerens* | Cloacal swab | Wild | Baja, Mexico |
| 12 | Crotalus cerastes | Sidewinder |  | Cloacal swab | Wild | Baja, Mexico |
| 13 | Pituophis catenifer | Gopher snake |  | Cloacal swab | Wild | Baja, Mexico |
| 14 | Chironius monticola | Vine snake | *Trichomitus batrachorum* | Cloacal swab | Wild | Colombia |
| 15 | Anolis fuscoauratus | Slender anole |  | Cloacal swab | Wild | Colombia |
| 16 | Gonatodes concinnatus | O'Shaughnessy Gecko |  | Cloacal swab | Wild | Colombia |
| 17 | Potamites ecleopus | Common stream lizard |  | Cloacal swab | Wild | Colombia |
| 18 | Paleosuchus trigonatus | Smooth-fronted caiman |  | Cloacal swab | Wild | Guyana |
| 19 | Platemys planicephala | Twist-necked turtle |  | Cloacal swab | Wild | Guyana |
| 20 | Chironius scurrulus | Smooth machete savane |  | Cloacal swab | Wild | Guyana |
| 21 | Corallus caninus | Emerald tree boa |  | Cloacal swab | Wild | Guyana |
| 22 | Corallus hortulanus | Amazon tree boa |  | Cloacal swab | Wild | Guyana |
| 23 | Uranoscodon superciliosus | Diving lizard |  | Cloacal swab | Wild | Guyana |
| 24 | Immantudes temnuissimus | Yucatan blunthead snake |  | Cloacal swab | Wild | Guyana |
| 25 | Bothrops asper | Lancehead pit viper |  | Cloacal swab | Wild | Guyana |
| 26 | Amphibaena fuliginosa | Speckled worm lizard |  | Cloacal swab | Wild | Guyana |
| 27 | Echis pyramidum | Carpet viper | *Hypotrichomonas acosta* | Cloacal swab | Wild | Ethiopia |
| 28 | Panaspis sp. nov. | Lidless skink | *Monocercomonas colubrorum* | Cloacal swab | Wild | Ethiopia |
| 29 | Chamaeleo dilepis | Flap-necked chameleon |  | Cloacal swab | Wild | Ethiopia |
| 30 | Hemidactylus isolepis | scaly leaf-toed gecko |  | Cloacal swab | Wild | Ethiopia |
| 31 | Lygodactylus keniensis | Parker's dwarf gecko |  | Cloacal swab | Wild | Ethiopia |
| 32 | Sitrurus miliarius | Rough green snake | *Monocercomonas colubrorum* | Cloacal swab | Wild | Florida, USA |
| 33 | Micrurus fulvis | Eastern coral snake | *Hypotrichomonas acosta* | Cloacal swab | Wild | Florida, USA |
| 34 | Pantherophis guttatus | Corn snake |  | Cloacal swab | Wild | Florida, USA |
| 35 | Thamnophis sirtalis | Garter snake |  | Cloacal swab | Wild | Florida, USA |
| 36 | Nerodia fasciata | Banded water snake |  | Cloacal swab | Wild | Florida, USA |
| 37 | Pantherophis obsoletus | Western ratsnake |  | Cloacal swab | Wild | Florida, USA |
| 38 | Ophisaurus attenuatus | Slender glass lizard |  | Cloacal swab | Wild | Florida, USA |
| 39 | Diadophis punctatus | Ringneck snake |  | Cloacal swab | Wild | Florida, USA |
| 40 | Coluber constrictor | Eastern racer |  | Cloacal swab | Wild | Florida, USA |
| 41 | Crotalus adamanteus | Eastern diamondback rattlesnake |  | Cloacal swab | Wild | Florida, USA |
| 42 | Opheodrys aestivus | Rough green snake |  | Cloacal swab | Wild | Florida, USA |

We identified 6 parabasalid species in these samples, but intriguingly only 4 species (*Hypotrichomonas acosta, Trichomitus batrachorum, Monocercomonas colubrorum*, and *Hexamastix coercens*) have been previously found in reptile microbiomes. The other two species were both novel species of the genus *Tritrichomonas* and were identified in rodent-eating snakes. These species phylogenetically clustered with the murine-associated protists *Tritrichomonas musculis* and *Tritrichomonas casperi*^15^, and thus are likely non-viable protist DNA from the intestines of rodents consumed by the snakes (Figure 1B). Of the remaining 4 reptile-associated parabasalid species identified in the wild reptiles, all but *H. coercens* were found in at least one captive reptile. To confirm that the reptile-associated parabasalid DNA sequences represented resident microbes, we isolated protists from the fresh stool of a domestic tortoise (*Testudo horsfieldii*) and identified motile *Trichomitus batrachorum* trophozoites that matched the ITS sequence from the unpurified stool DNA sample. Thus, the substantial overlap in parabasalid species between wild and captive reptiles and the high frequency of colonization in captive reptiles suggests that captivity does not have a detrimental impact on either parabasalid diversity or frequency in reptiles.

This survey also revealed a surprisingly low level of diversity among reptile-associated parabasalids. Only 4 reptile-associated species were identified, despite the broad host and geographical ranges sampled (Figure 1A, Table 1). Particularly striking was that 2 new rodent-associated parabasalids were discovered in the guts of rodent-eating snakes, yet no new reptile-associated protists were identified despite only surveying reptiles in this study. These data suggest that reptile-associated parabasalids have lower diversity than their mammal-associated counterparts. Supporting this hypothesis was the presence of *T. batrachorum* in both the captive tortoises (*Testudo horsfieldii, Testudo graeca*) in California and the wild vine snake (*Oxybelis fulgidus*) in Colombia. The tortoises sampled here are strict herbivores, whereas vine snakes are strict carnivores, and thus *T. batrachorum* appears adapted to hosts with diverse dietary patterns. This is particularly surprising because mammal-associated intestinal parabasalids have more specialized host ranges, suggesting that colonization of reptiles may require increased flexibility. Possible explanations for this phenomenon include less complex social structures than mammals, low incidence of vivipary, and rarity of parental involvement with offspring.

Next, we sought to understand how the drastic temperature fluctuations associated with climate change would affect the protists and their interactions with reptile hosts. Ectotherm-associated microbes, unlike microbes in the guts of endotherms, are exposed to cold temperatures *in vivo*, and thus we chose to look at adaptations to this temperature extreme. To look for common adaptations of diverse reptile-associated parabasalids, we cultured *T. batrachorum*, a member of class *Trichomitea, and M. colubrorum*, a member of class *Tritrichomonadea* (Figure 2A). We compared these protists to a human-associated species, *Pentatrichomonas hominis*, which colonizes endothermic mammals (Figure 2A). We then tested whether human or reptile-associated species would survive cold temperature by growing the three protists at 12°C overnight. As expected, this cold exposure was lethal for *P. hominis*, but the reptile-associated species survived and even grew at 12°C (Figure 2A, B). These data support the idea that reptile associated parabasalids have adapted to withstand substantial temperature fluctuations, as expected given their ectothermic hosts.

**Figure 2:**
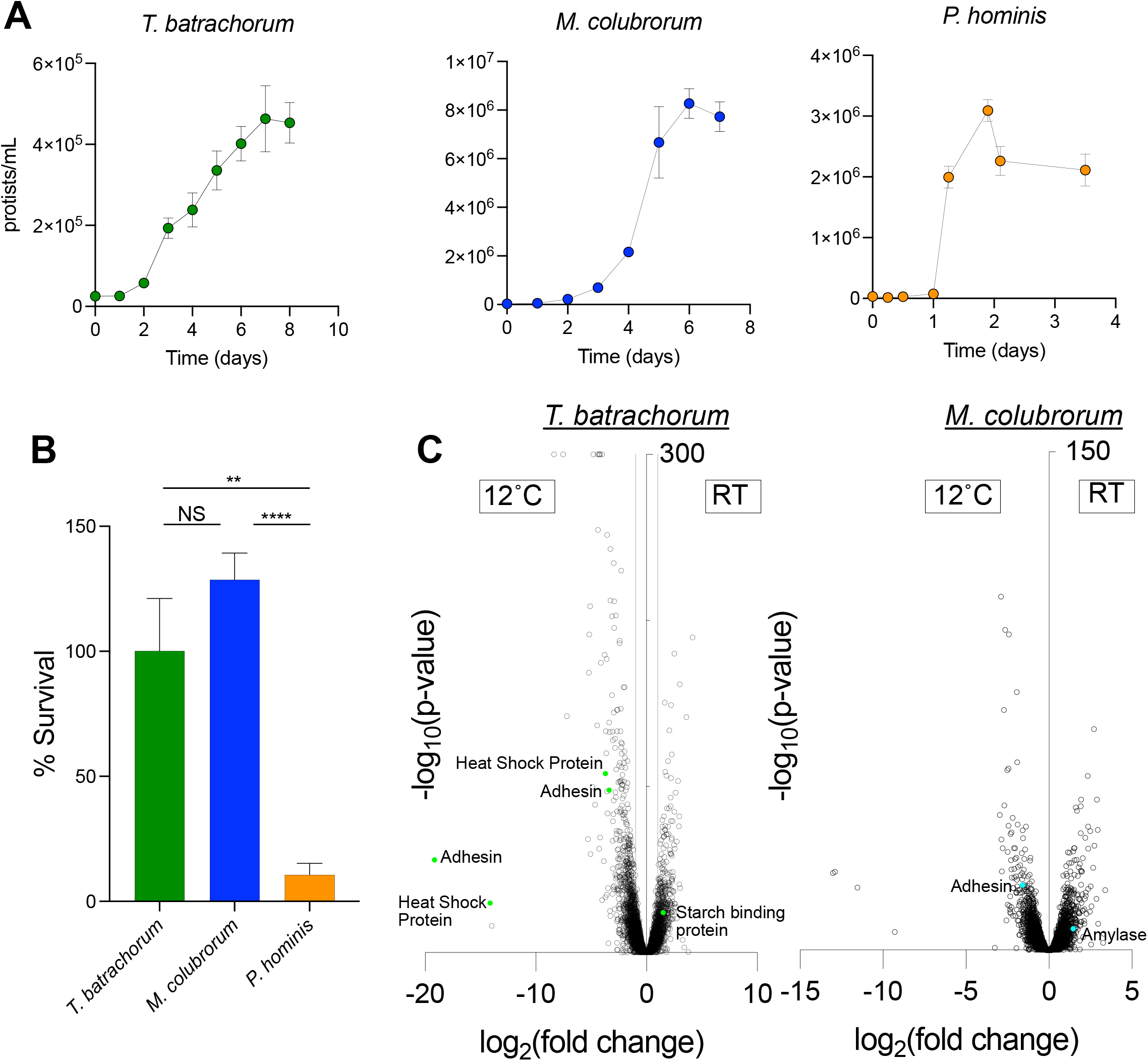
Reptile-associated parabasalids survive cold temperature and dramatically remodel their transcriptional profiles. (A) Growth curves of *T. batrachorum, M. colubrorum*, and *P. hominis* at their ideal growth temperatures (room temperature or 37°respectively). (B) Survival of the three protist species during overnight culture at 12°C. (C) Transcriptomic changes in *T. batrachorum* (left) and *M. colubrorum* (right) during growth at 12°C. Transcripts with increased expression during cold exposure are shown on the left. Select differentially expressed genes are colored (*T. batrachorum*, green; *M. colubrorum*, blue) and annotated with the gene function. NS not significant, **p<0.01, ****p<0.0001.

To determine whether cold exposure may affect the interactions of *T. batrachorum* and *M. colubrorum* with their hosts, we performed RNA-Sequencing and *de novo* transcriptome assembly on each protist during growth at either room temperature or 12°C. Cold exposure resulted in drastic transcriptional responses in both protists, as 1,808 (14.9% of detected transcripts) and 977 (5.2% of detected transcripts) genes were differentially expressed between the two temperatures in *T. batrachorum* and *M. colubrorum*, respectively (Figure 2C).

Strikingly, the most highly upregulated gene in *T. batrachorum* during growth at cold temperature encodes an adhesin protein, which was expressed more than 500,000-fold more highly at 12°C. An adhesin was also among the highly upregulated genes during cold exposure in *M. colubrorum*, indicating that this adaptation is common across reptile-associated parabasalids. These protists may adhere to the intestinal epithelium during cold exposure by upregulating adhesion proteins. This switch would be predicted to drastically change host-microbe interactions, as adhesion to the epithelium can trigger intestinal immune pathways^16^.

Genes involved in utilizing starch, a prominent plant polysaccharide, are downregulated during cold exposure in both protists. This suggests that the protists adapt to a predicted lack of plant material entering their reptilian host’s digestive tracts during periods of cold exposure. Importantly, we recently demonstrated that murine-associated parabasalids switch from fermenting dietary fiber to the host’s intestinal mucus glycans during fiber deprivation^15^. Thus, the downregulation of plant-polysaccharide utilization genes in reptile parabasalids suggests that these protists rely on the mucus glycans lining the epithelium during cold exposure. Importantly, the metabolic switch from plant polysaccharide utilization to host mucus glycans by the bacterial microbiota in mice erodes the intestinal mucus barrier and increases susceptibility to pathogens^17,18^. Altogether, these transcriptomic results suggest that parabasalid protists may respond to cold temperatures by adhering to the mucosa and fermenting host-derived carbohydrates, possibly to preempt the predicted decrease in activity and dietary intake by the reptilian host during colder temperatures. Importantly, these transcriptional changes by the protists are predicted to have detrimental effects on host health.

## Discussion

Parabasalid protists are increasingly recognized as part of animal microbiomes with significant influences on the health and immune function of their hosts. This study presents the first survey of these microbes in wild reptiles sampled across three continents and a diverse collection of captive reptiles. We found that parabasalid diversity is unusually low in reptiles compared to their mammalian counterparts and that reptile-associated protists have unprecedented flexibility in host range, suggestive of adaptations to the asocial lifestyle commonly found in reptiles. Our results also demonstrate that although captivity does not affect the diversity or prevalence of parabasalids in reptile microbiomes, the extreme weather patterns caused by climate change may dramatically affect the interactions of these microbes with their hosts. Specifically, *T. batrachorum* and *M. colubrorum* upregulate potentially detrimental genes during cold exposure, suggesting that extreme weather conditions may force commensal parabasalids to adopt pathobiont properties. Because the microbiota, including microeukaryotes like protists, critically impacts animal health, conservation efforts will be aided by understanding how captivity and environmental factors like climate change shape the composition and interactions of these microbes with vulnerable animal species.

## Methods

### Cloacal swabs and stool collection

Cloacal swabs have been shown to be an effective proxy for collecting the diversity of bacteria present in the GI tract of reptiles^19,20^. Swabs were collected using established methods for sampling reptile cloacal microbiomes^19,21^. Briefly, animals were either captured by hand or via pitfall traps according to approved protocols during biodiversity survey expeditions undertaken by TJC from 2010–2020. Once restrained, the exterior of the cloaca was cleaned with alcohol swabs or 95% ETOH, then a sterile nylon tipped swab was inserted into the cloaca and rotated 10 times, taking care not to penetrate into the large intestine. Swabs were then placed either in individual empty 1.5 ml cryovials and immediately frozen in liquid nitrogen^19^; or placed in 1.5ml cryovials containing 750 ul Xpedition™ RNA/DNA Shield (Zymo Research Products) and stored at ambient temperature before transportation to the laboratory where they were subsequently stored at -20°C until DNA extraction^22^. For captive reptiles, fresh stool samples were collected and frozen immediately.

### DNA extraction and PCR

DNA was extracted from cloacal swab and stool samples using the Powersoil Pro DNA Extraction kit (Qiagen) according to manufacturer instructions. Samples were screened for the presence of parabasalid protists by amplifying DNA using custom pan-parabasalid primers, which we designed to bind to regions of the internally transcribed spacer (ITS) that are highly conserved among the parabasalid lineage (panParabasalid-F 5’-CCACGGGTAGCAGCA-3’, panParabasalid-R 5’-GGCAGGGACGTATTCAA-3’). These primers amplify an approximately 1.1kb ITS amplicon. DNA was amplified using the Accustart II Geltrack Supermix (Quantabio) using touchdown PCR, with a starting annealing temperature of 72°C dropping to 65°C over 15 cycles, followed by 35 cycles with an annealing temperature of 65°C. A 1-minute extension at 72°C was used for both stages. The presence of parabasalid DNA in the samples was initially assessed by agarose gel confirmation, followed by verification and species identification using Sanger Sequencing (Molecular Cloning Lab). Phylogenetic trees were created using Phylogeny.fr^23^ based on ITS sequences, using default parameters.

### Isolation of *Trichomitus batrachorum* and protist culture

*Trichomitus batrachorum* was isolated from fresh stool of a captive *Testudo horsefieldii* tortoise. A fresh stool sample was washed 3 times in sterile phosphate buffered saline (PBS), then resuspended in 40% percoll. This slurry was then underlaid with 80% percoll and centrifuged at 1000xg for 10min with no brake. The interphase, containing *T. batrachorum*, was washed twice with sterile PBS and then resuspended in growth medium. Protists were then grown in an anaerobic chamber at room temperature (Coy Laboratory Products). *Monocercomonas colubrorum* strain Ns-1PRR and *Pentatrichomonas hominis* strain Hs-3:NIH were obtained from ATCC. Both reptile-associated protists were grown at room temperature, whereas *P. hominis* was grown at 37°C. For growth curves and survival tests, protists were counted using a hemocytometer.

### Cold exposure survival assay

*P. hominis, T. batrachorum*, and *M. colubrorum* were cultured in triplicate at their respective normal growth temperatures until they reached mid-log phase. Each culture was then divided in two, with one culture continuing growth at normal temperature and one being placed at 12°C. Protists were cultured overnight at the two temperatures, and then viable protists were counted in each condition on a hemocytometer.

### RNA Sequencing and *de novo* transcriptome assembly

RNA was isolated from mid-log phase protists grown at normal growth temperature or 12°C using the Direct-zol RNA Miniprep kit (Zymo). RNA libraries were generated using the KAPA Stranded mRNA Sequencing Kit (Roche), with poly-dT enrichment of mRNA. Samples were sequenced on a HiSeq (Illumina) using 2×150 read lengths. Because no genome sequences exist for *T. batrachorum* or *M. colubrorum, de novo* transcriptome assembly was performed using Trinity^24^. Transcript abundances were obtained using kallisto^25^ and differential expression analysis was performed using DESeq2^26^.

## Acknowledgements

Animals were handled under approved IACUC protocols (U of Mississippi SOP13-04; Florida State University #1836). We thank staff of the Steinhart Aquarium for their assistance in collecting stool samples. In addition, we thank Norm Cyr for his help in figure construction and Shelly Silverstein and Stanley Amieva for their generous donation of samples. TJC would like to thank the Ethiopian Wildlife Authority, Florida Fish and Wildlife Conservation Commission, Mexican SEMANART and National Commission of Natural Protected Areas, the Environmental Protection Agency of Guyana, and the Colombian Autoridad Nacional De Licencias Ambientales for granting collection and export permits. We also thank staff at the Stanford Functional Genomics Facility for their expertise in RNA-Sequencing library preparation. ERG is supported by the Stanford Pediatric IBD and Celiac Disease Research Program and the Stanford Maternal & Child Health Research Institute. Funding was provided to TJC via a J. William Fulbright Fellowship, the National Science Foundation (DEB 1501711) and the University of Mississippi Graduate College. Funding support for M.R.H. was provided by the National Institutes of Health (NIH) R01DK128292 and R21AI171222.

## Declaration of interests

The authors declare no relevant competing interests.

## Notes

### Competing Interest Statement

The authors have declared no competing interest.

## References

1. Rosenblatt, J. S. Outline of the evolution of behavioral and nonbehavioral patterns of parental care among the vertebrates: Critical characteristics of mammalian and avian parental behavior. Scand. J. Psychol. 44, 265–271 (2003).

2. Moeller, A. H. et al. The Lizard Gut Microbiome Changes with Temperature and Is Associated with Heat Tolerance. Appl. Environ. Microbiol. 86, (2020).

3. Whitfield, S. M. et al. Amphibian and reptile declines over 35 years at La Selva, Costa Rica. Proc. Natl. Acad. Sci. 104, 8352–8356 (2007).

4. Sinervo, B. et al. Erosion of Lizard Diversity by Climate Change and Altered Thermal Niches. Science 328, 894–899 (2010).

5. Redford, K. H., Segre, J. A., Salafsky, N., Rio, C. M. del & McAloose, D. Conservation and the Microbiome. Conserv. Biol. 26, 195–197 (2012).

6. Kohl, K. D. et al. Gut microbial ecology of lizards: insights into diversity in the wild, effects of captivity, variation across gut regions and transmission. Mol. Ecol. 26, 1175–1189 (2017).

7. Howitt, M. R. et al. Tuft cells, taste-chemosensory cells, orchestrate parasite type 2 immunity in the gut. Science 351, 1329–1333 (2016).

8. Chudnovskiy, A. et al. Host-Protozoan Interactions Protect from Mucosal Infections through Activation of the Inflammasome. Cell 167, 444–456.e14 (2016).

9. Schneider, C. et al. A Metabolite-Triggered Tuft Cell-ILC2 Circuit Drives Small Intestinal Remodeling. Cell 174, 271–284.e14 (2018).

10. Escalante, N. K. et al. The common mouse protozoa Tritrichomonas muris alters mucosal T cell homeostasis and colitis susceptibility. J. Exp. Med. 213, 2841–2850 (2016).

11. Cleveland, L. R. THE EFFECTS OF OXYGENATION AND STARVATION ON THE SYMBIOSIS BETWEEN THE TERMITE, TERMOPSIS, AND ITS INTESTINAL FLAGELLATES. Biol. Bull. 48, 309-[326]-1 (1925).

12. Hampl, V., Cepicka, I., Flegr, J., Tachezy, J. & Kulda, J. Morphological and Molecular Diversity of the Monocercomonadid Genera Monocercomonas, Hexamastix, and Honigbergiella gen. nov. Protist 158, 365–383 (2007).

13. Dimasuay, K. G. B. & Rivera, W. L. Molecular characterization of trichomonads isolated from animal hosts in the Philippines. Vet. Parasitol. 196, 289–295 (2013).

14. Céza, V., Pánek, T., Smejkalová, P. & Čepička, I. Molecular and morphological diversity of the genus Hypotrichomonas (Parabasalia: Hypotrichomonadida), with descriptions of six new species. Eur. J. Protistol. 51, 158–172 (2015).

15. Gerrick, E. R. et al. Metabolic diversity in commensal protists regulates intestinal immunity and trans-kingdom competition. http://biorxiv.org/lookup/doi/10.1101/2022.08.26.505490 (2022) xdoi:10.1101/2022.08.26.505490.

16. Atarashi, K. et al. Th17 Cell Induction by Adhesion of Microbes to Intestinal Epithelial Cells. Cell 163, 367–380 (2015).

17. Sonnenburg, E. D. & Sonnenburg, J. L. Starving our Microbial Self: The Deleterious Consequences of a Diet Deficient in Microbiota-Accessible Carbohydrates. Cell Metab. 20, 779–786 (2014).

18. Desai, M. S. et al. A Dietary Fiber-Deprived Gut Microbiota Degrades the Colonic Mucus Barrier and Enhances Pathogen Susceptibility. Cell 167, 1339–1353.e21 (2016).

19. Colston, T. J., Noonan, B. P. & Jackson, C. R. Phylogenetic Analysis of Bacterial Communities in Different Regions of the Gastrointestinal Tract of Agkistrodon piscivorus, the Cottonmouth Snake. Plos One 10, e0128793 (2015).

20. Eliades, S. J. et al. Gut microbial ecology of the Critically Endangered Fijian crested iguana (Brachylophus vitiensis): Effects of captivity status and host reintroduction on endogenous microbiomes. Ecol. Evol. 11, 4731–4743 (2021).

21. Eliades, S. J., Colston, T. J. & Siler, C. D. Gut microbial ecology of Philippine gekkonids: ecoevolutionary effects on microbiome compositions. FEMS Microbiol. Ecol. 98, 1–11 (2022).

22. Smith, S. N., Colston, T. J. & Siler, C. D. Venomous Snakes Reveal Ecological and Phylogenetic Factors Influencing Variation in Gut and Oral Microbiomes. Front. Microbiol. 12, 603 (2021).

23. Dereeper, A. et al. Phylogeny.fr: robust phylogenetic analysis for the non-specialist. Nucleic Acids Res. 36, W465–W469 (2008).

24. Grabherr, M. G. et al. Full-length transcriptome assembly from RNA-Seq data without a reference genome. Nat. Biotechnol. 29, 644–652 (2011).

25. Bray, N. L., Pimentel, H., Melsted, P. & Pachter, L. Near-optimal probabilistic RNA-seq quantification. Nat. Biotechnol. 34, 525–527 (2016).

26. Love, M. I., Huber, W. & Anders, S. Moderated estimation of fold change and dispersion for RNA-seq data with DESeq2. Genome Biol. 15, 550 (2014).

